# Ksp37 is a rapidly mobilized NK cell secretory protein that engages FGF2

**DOI:** 10.64898/2026.11.10.765584

**Authors:** Charlotte KJC Hermans, Kishan K Chudasama, Axel Berg-Larsen, Ellen M Haugsten, Merete Thune Wiiger, Alonso Sanchez, Silje Zandstra Krokeide, Agnieszka Graczyk-Bartnik, Marta Kłopotowska, Malgorzata Firczuk, Jørgen Wesche, Karl-Johan Malmberg, Aline Pfefferle

## Abstract

A key feature of natural killer (NK) cells is their preloaded arsenal of effector molecules, allowing them to rapidly respond upon activation. In this study, we characterized a previously unexplored secretory protein, termed killer-specific secretory protein of 37 kDa (Ksp37), which is highly abundant in differentiated CD56^dim^ NK cells. Ksp37 is rapidly secreted in a dose-dependent and JAK-1 dependent manner upon low levels of cytokine stimulation, making it a sensitive readout for NK cell activation. Following secretion, intracellular levels were rapidly replenished, preserving responsiveness to subsequent cytokine stimulation. We found that Ksp37 is stored in polarized intracellular compartments distinct from those containing canonical cytotoxic effector molecules, including granzyme B and perforin, which require stronger stimulation for release. Lastly, we identified Ksp37 as a novel binding partner of fibroblast growth factor 2 (FGF2). This opens new avenues of exploration for the role of NK cells in remodeling their extracellular environment by modulating the FGF2-FGFR1 axis, which plays a key role in tumor growth, angiogenesis and tissue repair.

## Introduction

Natural killer (NK) cells circulate the body in a primed state, ready to elicit their cytotoxic capacity upon encountering foreign, infected or malignant cells^1–3^. Their ability to rapidly respond as part of the innate immune system is dependent upon pre-loaded secretory granules containing cytotoxic effector molecules, such as granzyme B and perforin^4,5^. However, NK cells can also secrete numerous other effector molecules, such as chemokines (CCL3, CCL4, CCL5, XCL1) and cytokines (IFNy, TNFα), to modulate the surrounding environment by attracting and activating other immune cells^2,6,7^. While these effector molecules have been well-characterized, the function and secretion dynamics of the total secretome remains poorly understood^8,9^. One understudied secretory protein that is highly expressed in cytotoxic NK cells is killer-specific secretory protein of 37 kDa (Ksp37).

Ksp37, encoded by fibroblast growth factor binding protein 2 (*FGFBP2*), is the least well-characterized member of the FGFBP family which also contains *FGFBP1* and *FGFBP3*^10^. While *FGFBP1* is broadly expressed across various organs and *FGFBP3* is largely restricted to the brain, *FGFBP2* is solely expressed in NK and T cells^11,12^. Although *FGFBP2* is highly conserved across evolution, it is uniquely absent in the rodent genome^10,13^. Ksp37 was originally identified as a protein highly expressed in cytotoxic CD8 T cells compared to helper CD4 T cells, from which it received its name^11^. It is synthesized as a 28 kDa polypeptide which is heavily glycosylated post-transcriptionally and subsequently secreted as a 37 kDa protein^11^. All members of the FGFBP family contain a signaling peptide for secretion, a heparan-binding site and fibroblast growth factor (FGF)-binding sites, with binding of *FGFBP1* and *FGFBP3* to various FGF ligands having been reported^10,14,15^. Through binding to FGFs, FGFBPs can modulate availability of FGFs and in turn influence diverse physiological processes, such as cellular proliferation, differentiation and migration^10,16^.

FGF1 and FGF2 are potent mitogenic and angiogenic factors, which are secreted in a non-canonical fashion to ensure rapid mobilization in response to stress or injury^16,17^. They remain sequestered on the cell membrane by proximal heparan sulphate proteoglycan (HSPGs) and rely on FGFBPs for liberation and subsequent binding to FGF receptors (FGFR1-4)^10,17,18^. Elevated levels of Ksp37 have been detected in the sera of patients with an acute Epstein-Barr virus infection and in the bronchoalveolar lavage fluid of atopic asthma patients^11,19^. In the setting of cancer, *FGFBP2* expression in NK cells positively correlated with better prognosis in high-grade glioma, hormone receptor-positive breast cancer, hepatocellular carcinoma and epithelial ovarian carcinoma, while patients with autoimmune ankylosing spondylitis exhibited reduced expression of *FGFBP2*+ NK cells^20–24^. Despite these findings, and the prominent expression in NK cells, the biology of Ksp37 remains largely unexplored.

In this study we set out to perform a detailed characterization of the previously unexplored effector molecule Ksp37 in NK cells. Expression of Ksp37 correlated with differentiation, with functionally mature NK cells expressing high levels of Ksp37 which was expressed in a polarized fashion within the cell. Secretion of Ksp37 occurred in a highly regulated manner, dependent on the strength of cytokine signaling resulting in JAK1 activation. Intracellular stores of Ksp37 were rapidly replenished post-secretion, with sensitivity to cytokine-induced secretion being maintained over time. Lastly, we identified Ksp37 as a novel binder of FGF2, identifying a potential role of NK cells in modulating the FGF2-FGFR axis.

## Methods

### Cells

Buffycoats were obtained from anonymized healthy blood donors (Karolinska University Hospital, Sweden; Oslo University Hospital, Norway) with informed consent. The study was approved by the ethics committees in Sweden (Etikprövningsmyndigheten: Dnr. 2024-01771-02) and Norway (Regional etisk komité (REK): protocol no. 2018/2485). Peripheral blood mononuclear cells (PBMCs) were isolated by Ficoll-Paque density gradient centrifugation (Cytiva). NK cells were subsequently purified using magnetic-activated cell sorting (MACS, NK Cell Isolation kit, Miltenyi Biotec) and cultured in CellGenix® GMP Stem Cell Growth Medium (SCGM) supplemented with 10% human serum (Sigma-Aldrich) and 2 mM L-Glutamine (Gibco) unless otherwise stated. Medium was supplemented with IL-15 (0.1 to 10 ng/mL; R&D Systems), IL-2 (500 U/mL, Vecura), IL-12 (10 ng/mL; Peprotech), or IL-18 (100 ng/mL; Bio-Techne) to stimulate NK cells. Alternatively, NK cells were feeder-expanded using irradiated (100 Gy) K562-mIL-21-41BBL feeder cells (NK to feeder ratio 1:2) and 5 ng/mL IL-15 in a 24 well plate for 7 days. GolgiPlug^TM^ Protein Transport Inhibitor (1:2000; BD Biosciences) was added overnight to inhibit protein secretion. Cells were cultured for the indicated time periods at 37 °C and 5 % CO_2_.

### Bulk RNAseq dataset availability

To assess mRNA expression of *FGFBP2* within NK cell subsets, we used our previously published bulk RNA sequencing dataset containing sorted NK cell subsets from 10 healthy donors^25^.

### Inhibition of signaling pathways

To study signaling pathways involved in Ksp37 release, NK cells were cultured with various inhibitors overnight in the presence of 5 ng/mL IL-15: Ruxolitinib (JAK1/2 inhibitor, Invivogen; 1 μM), Fludarabine (STAT1 inhibitor, MedChemExpress; 5 μM), S3I-201 (STAT3 inhibitor, Sigma-Aldrich; 30 μM), Pimozide (STAT5 inhibitor, Merck Millipore; 7 μM), Rapamycin (mTORC1 inhibitor, Sigma-Aldrich; 50 nM), Torin 1 (mTORC1/2 inhibitor, APExBIO; 125 nM), PD98059 (MEK1/2 inhibitor, Merck; 50 μM), Wortmannin (PI3K inhibitor, Merck; 1 μM) or DMSO (Sigma-Aldrich) as negative control. Intracellular levels of Ksp37 were assessed after overnight incubation using flow cytometry.

### Flow cytometry

Viability and surface antigen staining (15 min) was performed in a 96 V-bottom plate, followed by fixation/permeabilization (20 min) and intracellular staining (30 min) at room temperature, using the following antibodies and reagents: CD16-BUV395 (3G8, 1:50), CD56-BUV737 (NCAM16.2, 1:25), Granzyme B-AF700 (GB11, 1:100), CD3-V500 (UCHT1, 1:100), Cytofix/Cytoperm^TM^ Fixation and Permeabilization Solution, Perm/Wash^TM^ buffer from BD Biosciences; CD57-PacificBlue (HNK-1, 1:50), Perforin-BV711 (δG9, 1:50), Ksp37-PE (TDA3, 1:100) from Biolegend; NGK2A-VioBright FITC (REA110, 1:100), NKG2C-APC (REA205, 1:50) from Miltenyi Biotec; CD158b1/b2-PE-Cy5.5 (GL183, 1:50), CD158a,h-PE-Cy7 (EB6B, 1:20) from Beckman Coulter; LIVE/DEAD Fixable Aqua Dead Cell Stain (1:100) from Invitrogen. Samples were acquired on a LSRFortessa (355, 405, 488, 561, 637 nm lasers; BD Biosciences) and analyzed using FlowJo version 10 (BD Biosciences).

### Cell sorting

Cells were stained for surface antigens and viability as previously described, followed by streptavidin staining (10 min) using the following antibodies and reagents: CD56-PE-Vio635 (REA196, 1:25), CD3-Biotin (REA613, 1:25) from Miltenyi Biotec; Streptavidin-BV711 (1:200) from BD Biosciences; LIVE/DEAD Fixable Aqua Dead Cell Stain (1:100) from Invitrogen. Sorting was performed on a Sony MA900 cell sorter (405, 488nm lasers; 100 μm nozzle) and cells were immediately stained for microscopy.

### Confocal microscopy

To investigate Ksp37 co-occurrence with effector molecules or vesicle markers, NK cells were cytokine (5 ng/mL IL-15, 4-5 days) or feeder-expanded respectively. Cells were harvested, seeded onto Tak^TM^ Cell and Tissue Adhesive (Corning) coated cover glasses (High precision 1.5H, Paul Marienfeld) and incubated at 37 °C (30 min) to adhere. Cells were subsequently fixed (Cytofix/Cytoperm^TM^ Fixation and Permeabilization Solution, BD Biosciences, 15 min), blocked (PBS + 10% goat serum (VWR), 1h), stained with primary antibodies (4h) and stained with secondary antibodies (30 min) combined with a membrane dye (Wheat Germ Agglutinin (WGA)-CF®640R (Biotium) at room temperature. Washing steps were performed using BD Perm/Wash^TM^ buffer (BD Biosciences) and samples were mounted in ProLong™ Diamond Antifade Mountant (Invitrogen). The following antibodies were used: Ksp37 (TDA3, 1:200), Perforin-AF488 (δG9, 1:200) from Biolegend; Rab5a (C8B1, 1:200), Rab7 (D95F2, 1:50), Rab8a (D22D8, 1:200), Rab9 (D52G8, 1:50) from Cell Signaling Technology; Rab18 (polyclonal, 1:400) from Sigma-Aldrich; MICALL1 (polyclonal, 1:150), LAMP1 (polyclonal, 1:200), Granulysin (EPR22110-101, 1:250) from Abcam; Granzyme B (496B, 1:100), Mouse IgG (polyclonal, 1:500), Rabbit IgG (polyclonal, 1:500), Rat IgG (polyclonal, 1:500) from Invitrogen. For staining with perforin-AF488, Ksp37 was directly conjugated with CoraLite® Plus 555 (FlexABLE 2.0 antibody labelling kit for Mouse IgG1 (Proteintech)).

### Image acquisition and analysis

Images were acquired on a Nikon spinning disk confocal running with the NIS Elements software and equipped with a Nikon Ti2 microscope, a BrixX HUB Ultra laser unit (Omikron-Laserage), a Kinetix camera (pixel size 6.5 µm, Photometrics) and a X-Light V3 spinning disk (50 µm pinholes, 250 µm interval, Crest Optics). The images were acquired with a 60×/1.4 oil objective. The excitation and emission wavelengths used to image included: Alexa Fluor Plus 405 (ex 405, em 438/24), Alexa Fluor Plus 488 (ex 488, em 515/30, Alexa Fluor Plus 555 (ex 555, em 593/40), and CF®640R (ex 638, em 685/40). To assess co-occurrence at higher resolution, images were acquired on a Nikon single point confocal equipped with NSPARC 3D-super-resolution detector. The images were acquired with a 100x/1.45 oil objective using the following excitation and emission filters: Alexa Fluor Plus 405 (ex 405, em 429-474), Alexa Fluor Plus 488 (ex 488, em 503-567, and CF®640R (ex 638, em 653-701). The imaging parameters were set to have no saturation in the regions of interest. To ensure a reproducible focus between samples, imaging was done right above the coverslip based on the WGA membrane stain. Z-stacks were taken to confirm that there were no chromatic aberrations.

Images were analyzed using a custom Fiji macro and script (ImageJ 1.54p; Java 21.0.7, 64-bit; Bio-Formats 8.5.0)^26^. NK cells were segmented on the membrane stain after background subtraction (rolling ball radius 60), Gaussian blur (sigma 2), and Huang auto-thresholding, with a minimum size of 20 µm². To quantify the Ksp37 expression, the mean Ksp37 signal of each cell was measured after background subtraction (rolling ball radius 8), and cells were classified as Ksp37 positive when this exceeded a fixed threshold, applied equally to all images. The percentage of Ksp37^+^ NK cells and the mean Ksp37 intensity of positive cells were calculated per image. To quantify co-occurrence with vesicle markers and effector molecules, cell segmentation was done as explained before. Next, double positive cells expressing both Ksp37 and the protein of interest to analyze its co-occurrence were classified using values set separately for each marker for background subtraction and thresholding. Next, co-occurrence was quantified per-cell using the Coloc 2 plugin (version 3.1.0, 100 Costes randomizations, PSF of 2.0) using Manders’ coefficient. For the generation of representative images, the image brightness and contrast were adjusted to allow detailed visualisation, with adjustments maintained across figure panels.

### Gene editing

Overnight IL-15 (10 ng/mL) stimulated NK cells were harvested and resuspended in P3 Primary Cell Nucleofector buffer (Lonza). CRISPR RNA (crRNA, IDT) and trans-activating crRNA (tracrRNA, IDT) were annealed at a 1:1 ratio at 95 °C for 5 min) to generate guide RNAs (gRNA). A ribonucleoprotein (RNP) complex of Cas9 protein (Alt-R™ S.p. Cas9 Nuclease V3, IDT) and three pooled gRNAs was formed at room temperature (10 min), before being combined with Alt-R™ Cas9 Electroporation Enhancer (IDT). In the presence of this RNP mixture, NK cells were electroporated using the pulse code CM 137 (Lonza 4D-Nucleofector). Cells were harvested and cultured as previously described for 7 days in the presence of 10 ng/mL IL-15. The following guide sequences purchased from IDT were used: *FGFBP2*_g1: 5’-GCGCCTTCACCATGCGTGCC-3’, *FGFBP2*_g2: 5’-CCCCCTGTACTCACACCAGT-3’, *FGFBP2*_g3: 5’-GGCAAGGTGCTGGAGAAGTC-3’, and the control gene *AAVS1*: 5’-CGGGGACACAGGAUCCCUGG-3’. Editing efficiency was assessed using flow cytometry.

### GST pulldown assays

FGF1 and FGF2 with an N-terminal GST tag were produced from pDest15(with rTEV-site)-GST-FGF1 (21-154) and pDest15(with rTEV-site)-GST-FGF2 (1-154)^27^ in *E.coli* Bl21 (DE3) pLyS and then purified on a Glutathione Sepharose 4 Fast Flow column. Recombinant GST protein was a generous gift from Coen Campsteijn (The Medical Faculty, University of Oslo).

[^35^S]-Methionine labelled FGFBP2 ([^35^S]-FGFBP2) was produced with pCMV6-Entry-FGFBP2 (RC206392, Origene) as template for *in vitro* transcription and translation using the TnT® Quick Coupled Transcription/Translation Systems with T7 polymerase (L1170, Promega) and [^35^S]-Methionine (NEG009A001MC, Promega).

For pulldown of GST-proteins with [^35^S]-FGFBP2, the recombinant GST/GST-FGF1/GST-FGF2 proteins were first allowed to bind to glutathione Sepharose 4 Fast Flow (Cytiva, 17-5132-01) in the presence of lysis buffer (0.1 M NaCl, 10 mM Na2HPO4, 1% Triton X-100, and 1 mM EDTA, pH 7.4) mixed 1:5 with PBS. Equimolar amounts of the GST proteins were used. Next, equal amounts of [^35^S]-FGFBP2 were added to each sample. The samples were rotated at 4°C for 1-4 hours, washed 3-5x in lysis buffer:PBS (1:5). The beads were then boiled with Laemmli loading buffer and samples were subjected to SDS-PAGE (4–20% Mini-PROTEAN® TGX™ Precast Protein Gel, #4561094, Bio-Rad). The gel was stained with Coomassie Blue to detect the recombinant GST-proteins and vacuum-dried. [^35^S]-FGFBP2 were detected with PharosFX Imager (Bio-Rad). Autoradiography images were quantified using Gel Analyzer in Fiji^26^.

### ELISA

An in-house ELISA was developed to quantify Ksp37 in cell supernatants and lysates. Supernatant was harvested from NK cells cultured overnight in serum-free SCGM (supplemented with 2 mM L-Glutamine), in the presence or absence of IL-15 (10 ng/mL) and/or GolgiPlug (1:2000) and cell pellets were lysed using Pierce RIPA lysis and Extraction Buffer (ThermoScientific), supplemented with cOmplete, EDTA-free Protease Inhibitor Cocktail (Merck).

A flat 96-well high binding plate (Corning) was coated with 2.5 μg/mL capture antibody (rabbit anti-human Ksp37, Proteintech, Cat No. 13254-1-AP) overnight at 4 °C. Unspecific binding sites were blocked with PBS supplemented with 0.05% Tween® 20 (PBS-T; Sigma-Aldrich) (2h, RT, gentle agitation) before NK cell lysate and supernatant samples were added (4 °C, overnight, gentle agitation). 4.3 μg/mL detection antibody (mouse anti-human Ksp37, BioLegend, clone TDA3) was added (1h, RT, gentle agitation) before incubating with HRP conjugated goat anti-mouse IgG (Agilent, Cat No. P0447, 1:3000 diluted) (1 h, RT, gentle agitation). As standard, Recombinant human Ksp37 was used (SinoBiological). Quantification was performed using the Ultra TMB-ELISA Substrate (Themo Scientific) and ELISA Stop Solution (Invitrogen), before measuring absorption at 450 nm using the xMark^TM^ Microplate Spectrophotometer (Bio-Rad).

### Statistical analysis

GraphPad Prism version 10.4.1 and 11.0.0 were used for data analysis and statistical testing. A Wilcoxon matched-pairs signed rank test or Mann-Whitney test was used for comparing two paired or unpaired groups respectively. A Friedman test or Kruskal-Wallis test was used to compare more than two paired or unpaired groups respectively, followed by Dunn’s post-hoc testing for multiple comparisons. A repeated measures two-way ANOVA with Geisser-Greenhouse correction, followed by Šídák post-hoc multiple comparison testing was used to compare grouped data with repeated measurements. A one-way ANOVA with Dunnett’s multiple comparisons test with a single pooled variance was performed for the FGF2-FGFP2 binding studies. p values: * <0.05, ** <0.01, *** <0.001, **** <0.0001.

## Code availability

The code used for the analysis of confocal microscopy data is available on GitHub and archived on Zenodo. Ksp37 expression quantification: GitHub, https://github.com/CharlotteKJCHermans/Ksp37-Expression-Quantification; Zenodo,

https://doi.org/10.5281/zenodo.23235614. Co-occurrence analysis: GitHub,

https://github.com/CharlotteKJCHermans/Ksp37-CoOccurrence-Quantification; Zenodo, https://doi.org/10.5281/zenodo.23474924.

## Results

### Ksp37 is abundant and expressed in a polarized pattern in cytotoxic NK cells

NK cell can be classified into distinct functional subsets based on their differentiation stage, as defined by their surface receptor repertoire (CD56, CD57, NKG2A, KIR)^28^. We had previously sorted five functionally distinct subsets from ten healthy donors for bulk RNA sequencing to generate a global expression profile across NK cell differentiation^25^. From these data we identified Ksp37, encoded by *FGFBP2*, as a highly expressed candidate that followed a clear maturation-dependent expression pattern (**Figure 1A-C**). While naïve CD56^bright^ NK cells expressed only low levels of *FGFBP2*, *FGFBP2* was one of the most highly expressed genes in cytotoxic CD56^dim^ NK cells, increasing in line with cytotoxic functional potential (**Figure 1A-C**).

**Figure 1.**
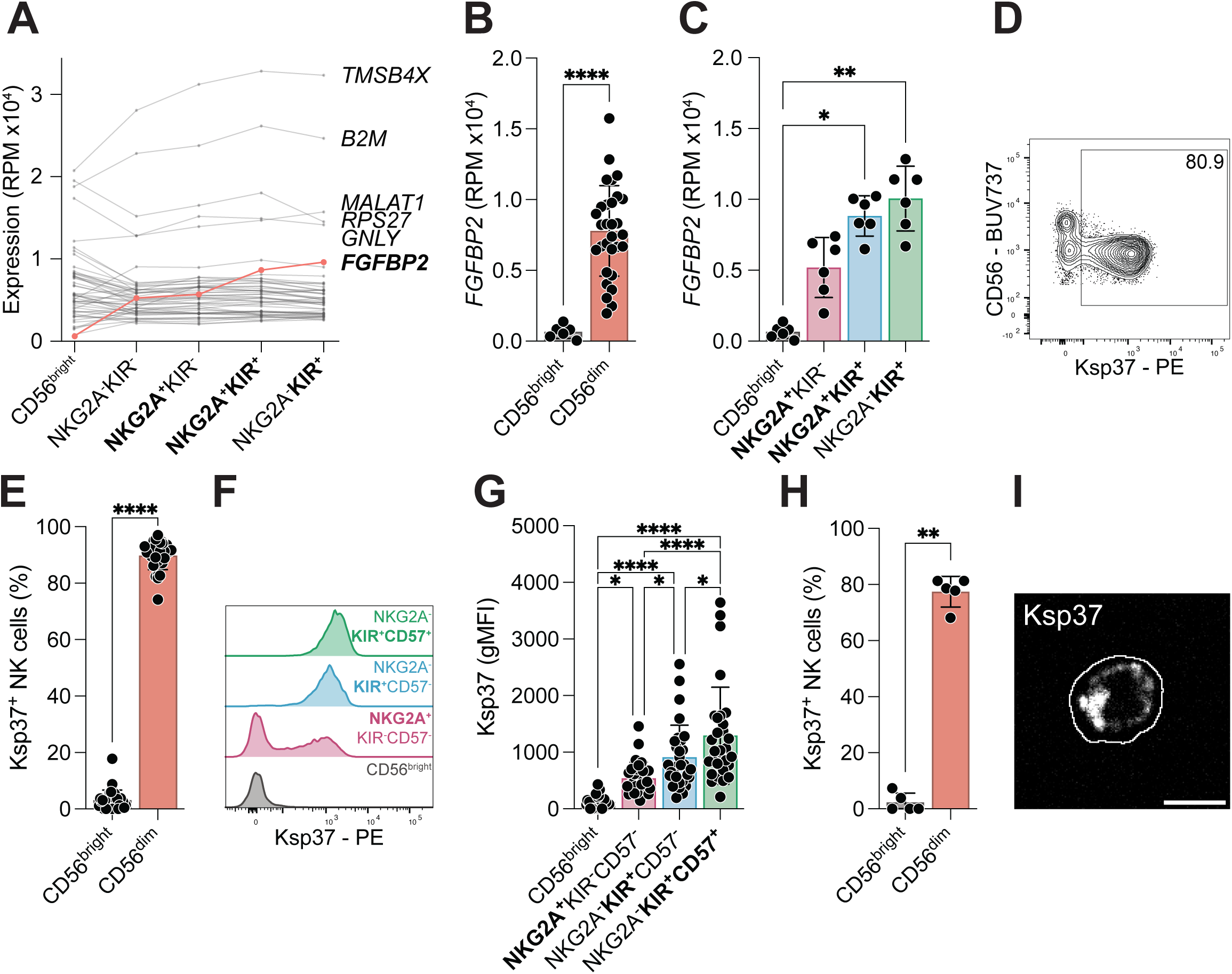
Ksp37 expression increases with NK cell maturation. (**A**) Expression of top 50 highest expressed genes in mature NKG2A^-^KIR^+^ NK cells among NK cell subsets. (**B**) mRNA expression of *FGFBP2* in CD56^bright^ and CD56^dim^ NK cells. (**C**) mRNA expression of *FGFBP2* in NK cell subsets. (**D**) Representative flow cytometry staining for Ksp37 in resting NK cells. (**E**) Percentage of Ksp37^+^ NK cells in CD56^bright^ and CD56^dim^ NK cells as quantified using flow cytometry. (**F**) Representative flow cytometry staining for Ksp37 expression in resting NK cell subsets as quantified using flow cytometry. (**G**) Ksp37 gMFI in NK cell subsets as quantified using flow cytometry. (**H**) Percentage of Ksp37^+^ cells in CD56^bright^ and CD56^dim^ NK cells as quantified using confocal microscopy. (**I**) Representative confocal image of Ksp37 (gray) in a CD56^dim^ NK cell. Cell outline is shown in white. Scale bar: 5 μm. Mann-Whitney tests were performed in (**B, H**). Friedman tests followed by Dunn’s multiple comparison tests were performed in (**C, G**). Wilcoxon test was performed in (**E**). Bars show the mean and error bars the standard deviation. *n* = 5 – 30 donors. \*\*\*\**p* < 0.0001; \*\**p* < 0.01; and \**p* < 0.05.

To verify expression at the protein level, we utilized flow cytometry to assess Ksp37 expression in the corresponding NK cell subsets (**Supplemental Figure 1A**). Confirming our findings at the mRNA level, Ksp37 was expressed in the large majority of CD56^dim^ NK cells, while only a small fraction of CD56^bright^ cells expressed the protein (**Figure 1 D-E**). Further stratification based on differentiation state showed a clear increase in both the frequency of Ksp37^+^ cells and the abundance of the protein in line with maturation (**Figure 1F-G**). Specificity of the antibody was confirmed by genetic deletion of *FGFBP2* in primary NK cells using CRISPR/Cas9 (**Supplemental Figure 1B-C**).

To decipher the expression pattern inside the cell, we sorted resting CD56^bright^ and CD56^dim^ NK cells and analyzed Ksp37 expression using confocal microscopy. Confirming flow cytometry results, CD56^bright^ NK cells expressed very low levels of the protein (**Figure 1H-I**). Within the sorted CD56^dim^ NK cells, Ksp37 was not only abundant but also appeared to be expressed in a polarized manner (**Figure 1H-I**). Hence, cytotoxic CD56^dim^ NK cells not only express higher levels of Ksp37, but also store it in a sequestered manner within the cell.

### NK cells rapidly secrete Ksp37 upon cytokine stimulation

Ksp37 is predicted to be a secretory protein, as it contains a common secretory signal in its C-terminal end and has been detected in sera from various patient groups^11,19,29^. We therefore set out to identify how secretion of the protein is triggered. Cytokines, in particular IL-15, are known to induce production and release of effector molecules and chemokines from NK cells. Overnight stimulation with IL-15 resulted in a significant decrease in Ksp37 expression, indicative of either secretion or protein degradation (**Figure 2A-B**). Brefeldin A (BfA) treatment prevented the IL-15-induced loss of intracellular Ksp37 (**Figure 2A-B**). Quantification of Ksp37 in NK cell culture supernatants and cell lysates supported secretion as the explanation for the decrease of intracellular Ksp37 after IL-15 stimulation (**Figure 2C-D**). This secretion in response to IL-15 stimulation was dose-dependent, as observed by a gradual decrease in Ksp37 expression with increasing doses of IL-15 (**Figure 2E-F**). More naïve NK cell subsets exhibited increased sensitivity to cytokine-mediated input compared to more mature subsets, which favor receptor-mediated input^7^. We could observe this IL-15 sensitivity at the subset level based on the amount of Ksp37 secreted by each subset in response to the same concentration of IL-15 (**Figure 2G-H**).

**Figure 2.**
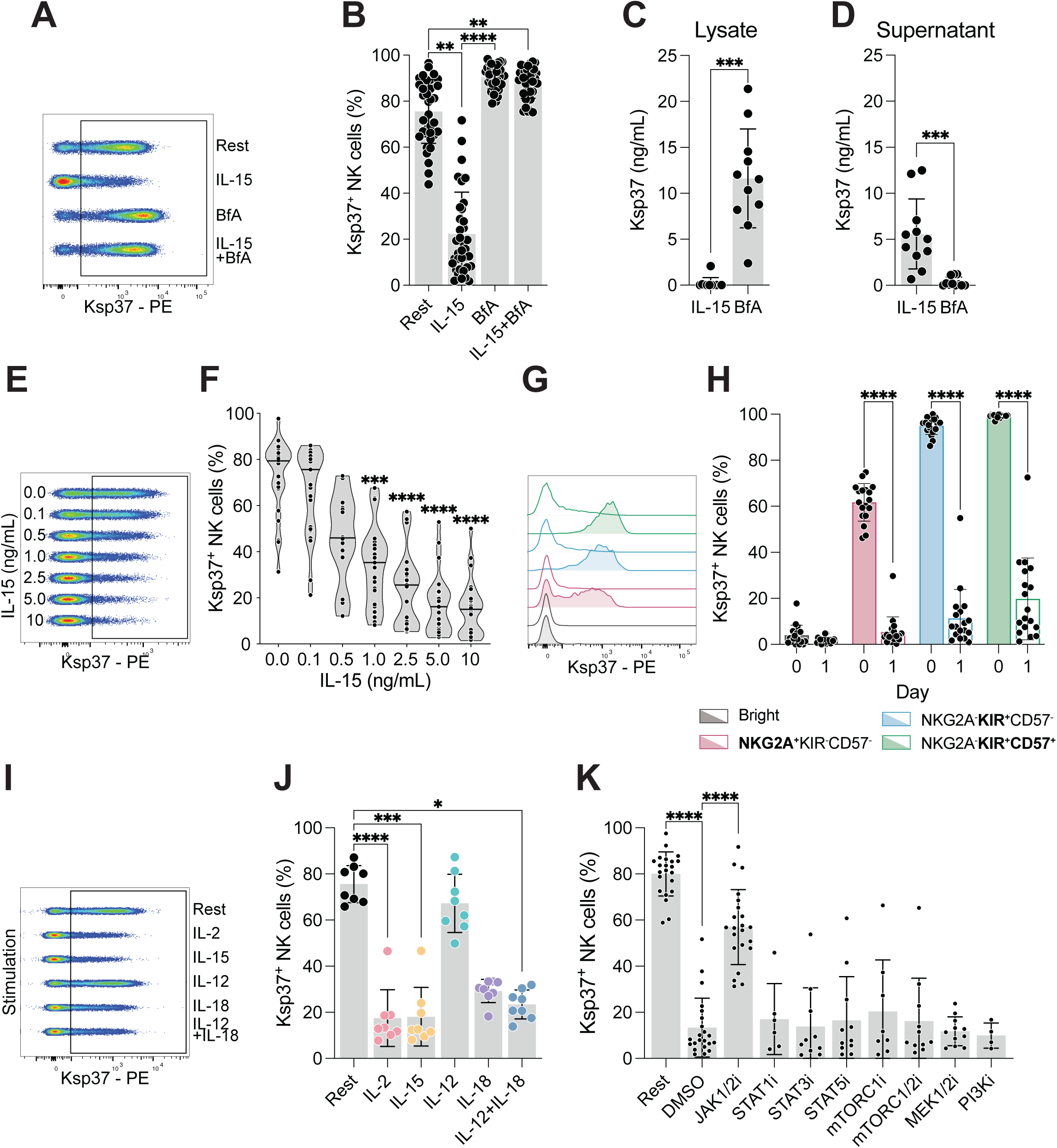
Rapid secretion of Ksp37 upon cytokine stimulation. (**A**) Representative flow cytometry staining showing Ksp37 expression in resting NK cells and in NK cells cultured with IL-15, brefeldin A (BfA), or both. (**B**) Percentage of Ksp37^+^ cells in resting NK cells and in NK cells cultured with IL-15, BfA, or both. (**C**) Concentration of Ksp37 in cell lysates from NK cells treated with either IL-15 or BfA. (**D**) Concentration of Ksp37 in supernatant from NK cells treated with either IL-15 or BfA. (**E**) Representative flow cytometry staining showing Ksp37 expression in NK cells treated with increasing concentrations of IL-15. (**F**) Percentage of Ksp37^+^ NK cells treated with increasing concentrations of IL-15. Asterisks indicate significance compared with untreated NK cells (no IL-15). (**G**) Representative flow cytometry staining showing Ksp37 expression in NK cell subsets at baseline (unfilled) and after overnight stimulation with 10 ng/mL IL-15 (filled). (**H**) Percentage of Ksp37^+^ cells in NK cell subsets at baseline (day 0, unfilled) and after overnight stimulation with 10 ng/mL IL-15 (day 1, filled). (**I**) Representative flow cytometry staining showing Ksp37 expression in resting NK cells and in NK cells cultured overnight with different cytokines (500 U/mL IL-2, 10 ng/mL IL-15, 10 ng/mL IL-12, 100 ng/mL IL-18). (**J**) Percentage of Ksp37^+^ NK cells upon rest or following overnight stimulation with different cytokines. (**K**) Percentage of Ksp37^+^ NK cells upon rest or with 5 ng/mL IL-15 in presence of indicated signaling pathway inhibitors (**I**) or DMSO vehicle control. JAK1/2i: 1 μM Ruxolitinib; STAT1i: 5 μM Fludarabine; STAT3i: 30 μM S3I-201; STAT5i: 7 μM Pimozide; mTORC1i: 50 nM Rapamycin; mTORC1/2i: 125 nM Torin 1; MEK1/2i: 50 μM PD98059; PI3Ki: 1 μM Wortmannin. Friedman tests followed by Dunn’s multiple comparison tests were performed in (**B, J**). Wilcoxon tests were performed in (**C, D**). Kruskal-Wallis tests followed by Dunn’s multiple comparison tests were performed in (**F, K**). Repeated measures two-way ANOVA with Geisser-Greenhouse corrected followed by Šídák’s multiple testing comparison test was performed in (**H**). Bars show the mean and error bars the standard deviation. *n* = 4 – 40 donors. \*\*\*\**p* < 0.0001; \*\*\**p* < 0.001; \*\**p* < 0.01; and \**p* < 0.05.

IL-15 plays an important role in NK cell development, proliferation and activation, however, other cytokines can also lead to NK cell activation, such as IL-2, IL-12 and IL-18, among others^30–32^. To understand if secretion of Ksp37 was unique to IL-15 or a more general response to cytokine stimulation, we repeated the overnight stimulation in the presence of IL-2, IL-12, IL-18 and IL-12/IL-18. IL-2 and IL-15 were the most potent inducers of Ksp37 secretion, while IL-18 and IL-12/IL-18 also induced significant amounts of Ksp37 secretion (**Figure 2I-J**). Interestingly, IL-12 stimulation alone could not induce secretion, while BfA treatment prevented secretion irrespective of stimulation (**Figure 2I-J**, **Supplemental Figure 2A-B**).

To further dissect the signaling pathway leading to Ksp37 secretion, we blocked various signaling molecules downstream of IL-2, IL-15, IL-12 and IL-18 signaling, including JAK1/JAK2, STAT1, STAT3, STAT5, mTORC1, mTORC1/2, MEK1/2 and PI3K (**Supplemental Figure 2C**). Secretion of Ksp37 was only significantly inhibited by blocking JAK1/2, indicating that JAK1 is likely required for cytokine-induced Ksp37 secretion (**Figure 2K, Supplemental Figure 2D**). Thus, secretion of Ksp37 by NK cells is a highly fine-tuned process in which the strength of cytokine signaling leading to JAK1 activation determines the amount of Ksp37 secreted.

### Ksp37 intracellular stores are rapidly replenished upon secretion

Secretion of Ksp37 occurs in a rapid, but controlled manner, resulting in intracellular stores being depleted after overnight cytokine activation. This raises the question of whether this release is a restricted event, or if intracellular stores are replenished to sustain secretion over prolonged activation periods. To decipher the dynamics of Ksp37 production and secretion, we performed longitudinal time-course experiments. After the initial overnight IL-15 stimulation, NK cells were kept in culture and monitored daily for intracellular Ksp37 expression. Without the addition of fresh cytokine addition, Ksp37 expression increased significantly over time, as extracellular IL-15 levels diminished due to cellular uptake (**Figure 3A-B**). However, sensitivity to subsequent cytokine stimulation was retained, as addition of fresh IL-15 (day 5, 7, 8) resulted in further secretion, as noted by diminished expression (day 6, 8, 9) (**Figure 3A-B**). This was confirmed by splitting the culture into two on day 5 and 8, with IL-15 only being added to one fraction. A significant decrease in intracellular Ksp37 levels, indicative of secretion, is noted < 24h after fresh cytokine addition (**Figure 3C-D**). Hence cytokine-induced activation induces both secretion but also production of fresh Ksp37 in NK cells to replenish intracellular stores.

**Figure 3.**
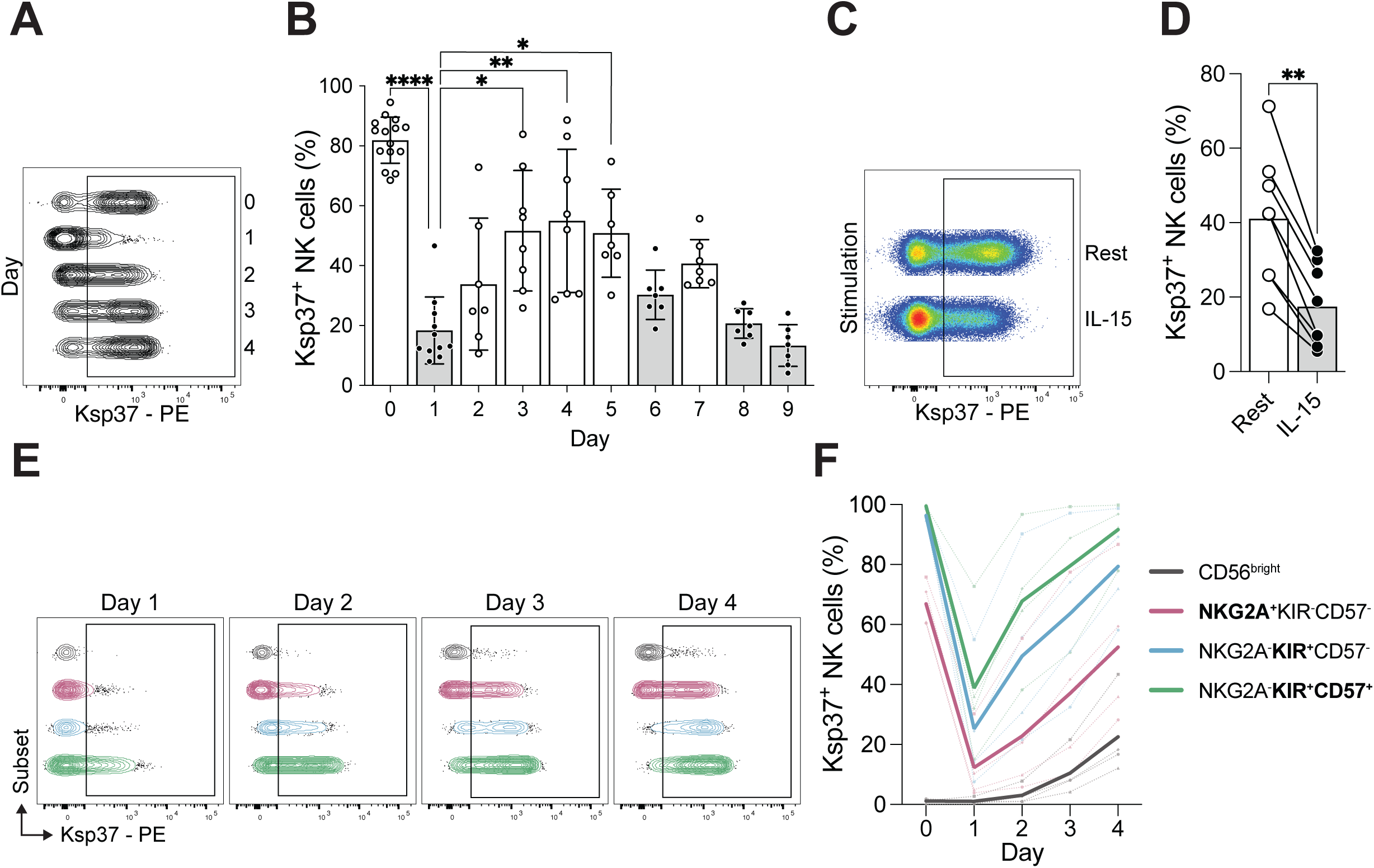
Post-stimulation reloading kinetics of Ksp37. (**A**) Representative flow cytometry staining showing Ksp37 expression prior to stimulation (Day 0) and on day 1-4 of stimulation with 5 ng/mL IL-15. (**B**) Percentage of Ksp37^+^ NK cells over time upon stimulation with 5 ng/mL IL-15. Bars highlighted in gray (day 1, 6, 8, 9) indicate the addition of new IL-15. (**C**) Representative flow cytometry staining showing Ksp37 expression on day 6 after overnight restimulation with fresh IL-15 or rest without fresh IL-15. (**D**) Percentage of Ksp37^+^ NK cells after overnight restimulation with fresh IL-15 or rest without fresh IL-15. Combined data from day 6 and 9. (**E**) Representative flow cytometry staining showing Ksp37 expression in NK cell subsets on day 1-4 of stimulation with 5 ng/mL IL-15. (**F**) Percentage of Ksp37^+^ cells within NK cell subsets before stimulation (day 0) and on days 1-4 of stimulation with 5 ng/mL IL-15. Solid lines indicate the mean and dashed lines indicate individual donors (*n* = 4). Kruskal-Wallis test followed by Dunn’s multiple comparison test was performed in (**B**). Wilcoxon test was performed in (**D**). Bars show the mean and error bars the standard deviation. *n* = 7 – 15 donors. \*\*\*\**p* < 0.0001; \*\**p* < 0.01; and \**p* < 0.05.

Cytokine-induced remodeling of surface receptor expression and effector molecules is well documented in NK cells^33,34^. Granzyme B and perforin are both rapidly upregulated in CD56^bright^ NK cells upon IL-15 stimulation, despite their secretory granules being largely devoid of these effector molecules at steady state^35^. We therefore wanted to explore if Ksp37 followed a similar trend or if expression was more restricted even post cytokine activation. Stratifying our time-course data into individual NK cell subsets, we observed a clear retention of the original expression pattern observed at steady state, with baseline levels of Ksp37 expression largely being recovered by day 4 of culture (**Figure 3E-F, Supplemental Figure 3**). Thus, NK cells readily secrete Ksp37 in a dose-dependent manner in response to cytokine-stimulation and rapidly replenished their intracellular stores to ensure available cellular stores for further secretion.

### Ksp37 localization is distinct from other known effector molecules

NK cells, upon activation, are efficient secretors of both pre-stored and newly synthesized effector molecules, cytokines and chemokines, allowing them to interact with surrounding cells and the larger environment^36,37^. To further characterize this novel secretory protein, we set out to identify how it is stored and secreted by NK cells using confocal microscopy. Feeder-expanded NK cells were used to capture a timepoint whereby NK cells were activated, expressed high levels of effector molecules and vesicular transport machinery (Rabs) required for secretion. Importantly, no fresh cytokines were added 48h prior to readout to allow for cellular stores of Ksp37 to be refilled. Ksp37 did not co-occur with LAMP1 (0.20), Perforin (0.17) nor Granzyme B (0.11), with some co-occurrence observed with Granulysin (0.35) (**Figure 4A-B**). As no significant co-occurrence was observed with known effector molecules nor lytic granules (LAMP1), we tested for co-occurrence with known markers for endocytic and transport vesicles, namely: Rab5a (early endosomes, incoming endocytic vesicles), Rab7 (late endosomes, transition to lysosomes), Rab18 (lipid droplets, endoplasmic reticulum), Rab9 (late endosomes, transport to TGN), Rab8a (post-Golgi exocytic transport vesicles, recycling endosomes), MICALL1 (tubular recycling endosomes) (**Figure 4C**). Co-occurrence was very low with all markers tested, with Rab7 scoring the highest (0.28) among these (**Figure 4D**). Hence, Ksp37 appears to be stored separately from other known effector molecules and is not restricted to a particular type of endocytic/trafficking vesicle, in line with its unique secretion dynamics.

**Figure 4.**
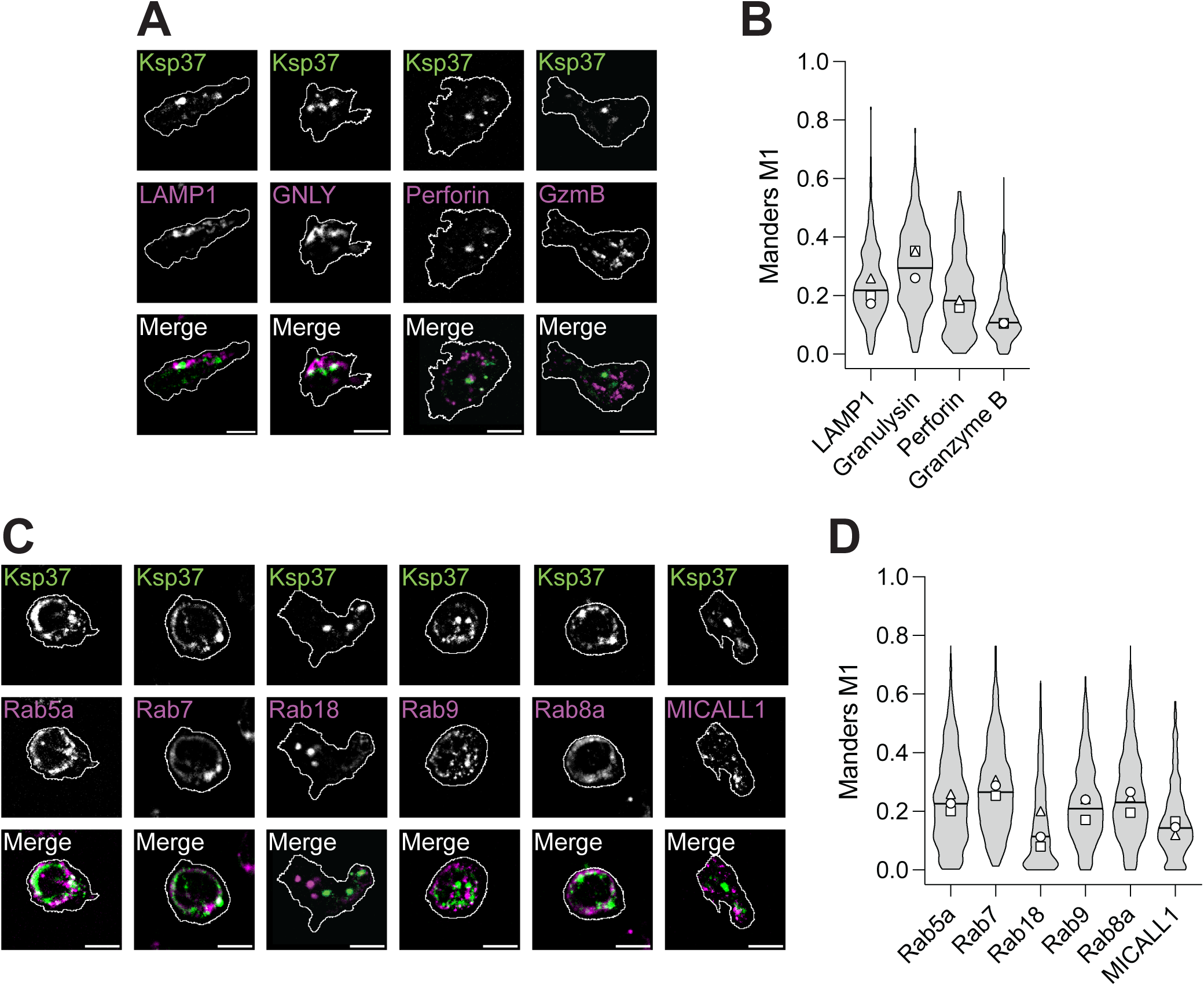
Ksp37 localizes distinct from other effect molecules. (**A**) Representative confocal images of NK cells co-stained for Ksp37 (green) and LAMP1, granulysin (GNLY), perforin, or granzyme B (GzmB) (magenta). Cell outlines are shown in white. Scale bar: 5 μm. (**B**) Co-occurrence of Ksp37 with LAMP1, granulysin, perforin, and granzyme B, quantified from confocal images as Manders’ coefficient M1 (fraction of Ksp37 signal overlapping with each effector molecule). Violin plots show all analyzed cells (≥ 69 per donor and condition); symbols indicate the median per donor (*n* = 2-3 donors). (**C**) Representative confocal images of NK cells co-stained for Ksp37 (green) and Rab5a, Rab7, Rab18, Rab9, Rab8a, and MICALL1 (magenta). Cell outlines are shown in white. Scale bar: 5 μm. (**D**) Co-occurrence of Ksp37 with Rab5a, Rab7, Rab18, Rab9, Rab8a, and MICALL1, quantified from confocal images as Manders’ coefficient M1 (fraction of Ksp37 signal overlapping with each effector molecule). Violin plots show all analyzed cells (≥ 132 per donor and condition); symbols indicate the median per donor (*n* = 3 donors). Optical resolution of 226 nm.

### Ksp37 is a binding partner of fibroblast growth factor 2 (FGF2)

High expression at steady state, dose dependent secretion in response to stimulation combined with replenishment of intracellular stores post secretion hints at Ksp37 having an important biological function. *FGFBP2*/Ksp37 is the least characterized member of the FGFBP family, which includes *FGFBP1* and *FGFBP3* which are known to bind FGF1 and/or FGF2, among others. Although *FGFBP2* has a predicted binding domain for FGF2 (**Figure 5A**), binding has not been shown to date. We used a rabbit reticulocyte system to express radioactively labelled Ksp37 and performed a pulldown assay using GST-tagged FGF1 and FGF2 to identify potential binding to Ksp37 (**Figure 5B**). A clear binding between FGF2 and Ksp37 was detected via autoradiography, while binding to FGF1 was negligible (**Figure 5C-D**). Hence, we have identified Ksp37 as a novel binding partner of FGF2, identifying a potential role of NK cells in modulating the FGF2-FGFRs axis.

**Figure 5.**
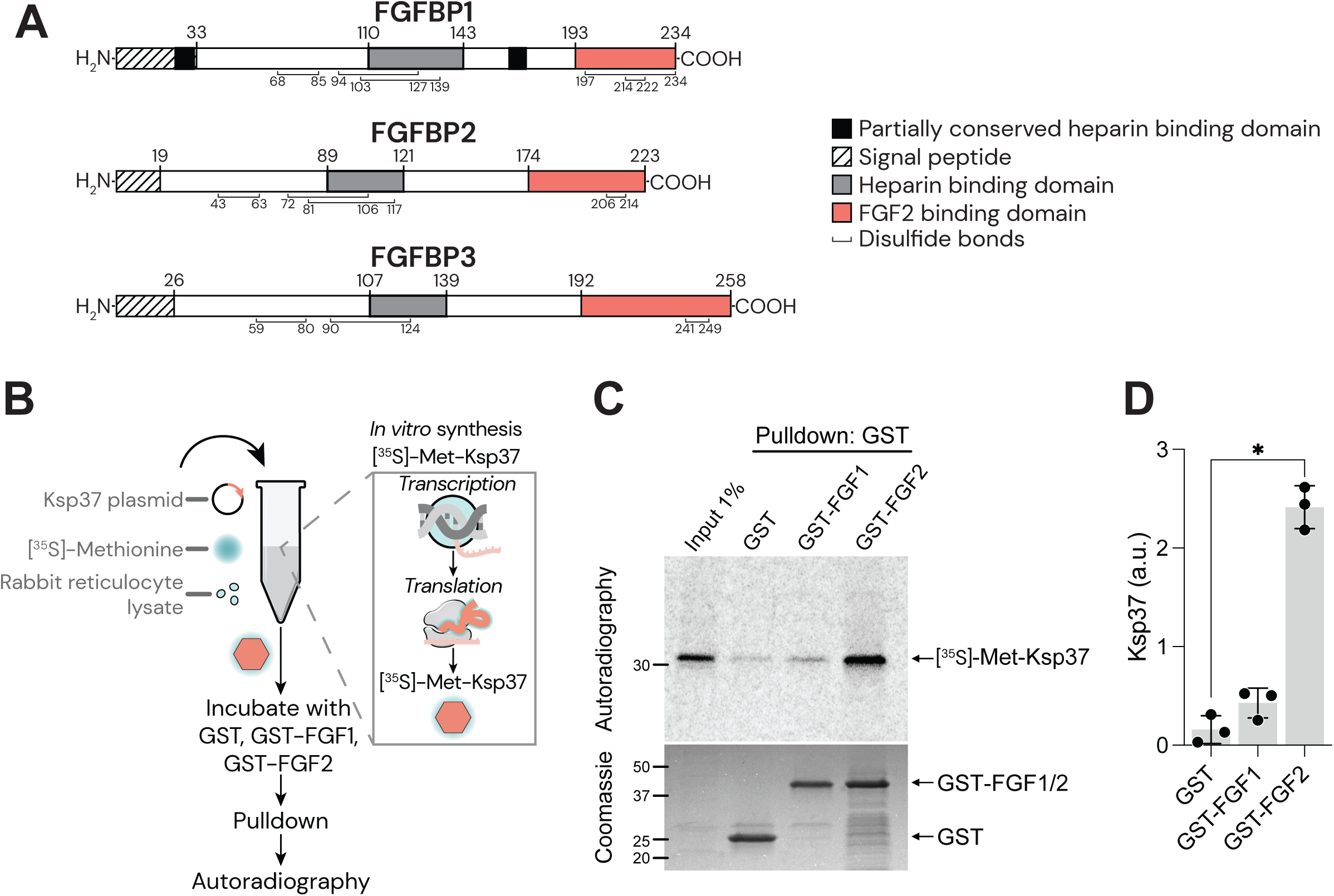
Ksp37 is a binding partner of FGF2. (**A**) Schematic of protein domains of FGFBP1-3. (**B**) Schematic of *in vitro* synthesis of [^35^S]—Methionine (Met)-Ksp37 and workflow for detection of binding to FGF1 and FGF2. (**C**) Representative image of autoradiography and Coomassie staining following pulldown of [^35^S]-Met-Ksp37 with GST, GST-FGF1, and GST-FGF2. (D) Quantification of [^35^S]-Met-Ksp37 binding to GST, GST-FGF1, and GST-FGF2. The bands corresponding to [^35^S]-Met-Ksp37 were quantified and normalized to mean of all for each experiment. Bars show the mean and error bars the standard deviation. Friedman test followed by Dunn’s multiple comparison test was performed in (**D**). *n* = 3 samples. \**p* < 0.05.

## Discussion

In this present study, we report a detailed characterization of an effector protein highly abundant in cytotoxic CD56^dim^ NK cells named Ksp37. The increased expression observed in more differentiated and functionally mature NK cells was intracellularly sequestered in a polarized manner, which was independent from other known effector molecules and not restricted to a specific endocytic vesicle. Secretion of Ksp37 was highly sensitive and fine-tuned to the level of cytokine-induced JAK1 activation. Upon secretion, intracellular stores of Ksp37 were efficiently restored to steady-state levels based on differentiation state. NK cells remained sensitive to further cytokine-induced Ksp37 secretion, highlighting the biological importance of this novel effector molecule. Lastly, we were able to demonstrate binding of Ksp37 to FGF2, opening new avenues of exploration to identify the functional role of the most abundant secretory protein in NK cells.

Litte is known about *FGFBP2* and its functional role, despite *FGFBP2* being commonly used in cytotoxic NK cell gene signatures due to its high expression^22,24,38^. Its restricted expression profile, contributing to the protein first being discovered, hints at a cytotoxicity-related role, as expression is restricted to T and NK cells, peaking in the more cytotoxic subsets. Further clues on its potential functional role can be delineated from its gene family, the fibroblast growth factor binding proteins (FGFBPs). Both *FGFBP1* and *FGFBP3* bind an array of FGFs, with FGFBP1 mobilizing locally stored FGF1/2 and FGFBP3 outcompeting heparan from binding to FGF2^14,15^. However, likely due to its more restricted expression compared to its family members, binding of *FGFBP2* to FGF2 had not previously been reported prior to this study in humans. Due to its potency, FGF2 is trapped in proximal heparan sulphate proteoglycan (HSPGs) structures on the surface of FGF2^+^ cells, requiring a binding protein for its release and subsequent binding to FGFR1^17^. Numerous ligands can bind FGFR1, with the downstream cellular effects being dependent on receptor fate due to ligand affinity^17^. One previously described ligand of low (micromolar) affinity is NCAM-1 (CD56) which is expressed on NK cells^39–45^. NCAM-1 binds FGFR1 as a monomer, leading to rapid recycling of the receptor and resulting in cellular migration^42,45–47^. The high (picomolar) affinity ligand FGF2, together with HSPG as a co-receptor, bind FGFR1 as a homodimer, resulting in downstream signaling inducing proliferation. The receptor complex is also internalized and degraded^46,48^. Another ligand binding FGFR1 as a homodimer, is galectin 3, which due to its low (micromolar) affinity, induces clustering of the receptor on the cell surface leading to cellular senescence^49^. It is therefore temping to speculate that CD56^bright^ NK cells, compared to CD56^dim^ NK cells would have distinct downstream effects on FGFR1^+^ cells in their surroundings, as the ratio of CD56 to Ksp37 inversely correlates. However, the binding strength of FGF2-Ksp37 to FGFR1 has not been determined. To identify the downstream effects of secreted Ksp37 by NK cells, the interplay between CD56-FGFR1 and Ksp37-FGF2-FGFR1 needs to be investigated further.

Secretion of Ksp37 is a highly sensitive measurement of NK cell activation by cytokine stimulation. Very low levels, 0.1 ng/mL of IL-15 given overnight, are already sufficient to induce secretion. Such low levels of IL-15 are insufficient to induce proliferation, degranulation or upregulation of other effector molecules such as granzyme B^50,51^. Hence, Ksp37 in the sera is indicative of NK cell activation and correlates with the magnitude of the response. Indeed, increased levels of Ksp37 have been detected in patient serum in the context of infection, cancer and autoimmune disease^12,19–21,23,24^. Ksp37 did not co-localize with key effector molecules, including perforin and granzyme B and it is unlikely to play a role in cellular cytotoxicity due its low threshold for release. A more likely role is remodeling of extracellular environment in inflammation or settings of elevated cytokine levels. Although not tested experimentally, we hypothesized that Ksp37 could contribute to remodeling of the uterine environment during pregnancy due to high levels of IL-15 and the importance of FGF2 during trophoblast invasion. Uterine NK cells are highly granular and have a secretory phenotype, but do not express *FGFBP2*, in line with their naïve-like differentiation state^52^. Hence secretion of Ksp37 in the uterus would be limited to peripheral blood-derived NK cells infiltrating the tissue, which are few in number.

Here we characterized a previously unexplored secretory molecule termed Ksp37, expression of which positively correlated with differentiation state. Rapid secretion in response to low levels of cytokine stimulation makes secreted Ksp37 a uniquely sensitive measurement of NK cell activation. Rapid replenishment of intracellular stores upon secretion, hints at the biological importance of this FGF2-binding novel secretory protein, with further studies required to pinpoint its exact functional role in the broader context of FGF2 signaling.

## Author contributions

CKJCH designed research, performed experiments, analyzed data, wrote the manuscript. KKC, AB-L, EMH designed research, performed experiments, analyzed data. MTW, AS, SZK, AG-B, MK performed experiments, analyzed data. MF, JW designed research, supervised the project. K-JM designed research, supervised the project, edited the manuscript. AP designed research, performed experiments, analyzed data, supervised the project, wrote the manuscript. All authors reviewed the manuscript.

## Acknowledgements

This study was in part performed at the Luminous Cell Imaging core facility/Nikon Center of Excellence, at Karolinska Institutet, Sweden, supported by the KI infrastructure board (1– 48/2024) and the Olle Engvist Foundation (235-0496). The work was further supported by the Research Council of Norway (275469, 237579, KJM), the Research Council of Norway through its Centres of Excellence scheme, project number 332727 (KJM) and FRIPRO 360808 (KJM), the Norwegian Cancer Society (190386, 223310, 321731 KJM), The South-Eastern Norway Regional Health Authority (2021-073, 2024-053, KJM), the Swedish Cancer Society (24 3917 Pj (KJM), 26 5114 IA JIA (AP)), the Swedish Children’s Cancer Society (PR2024-0148), Knut and Alice Wallenberg Foundation 2018.0106 (KJM), and the US National Cancer Institute (P01 CA111412 (KJM), P009500901 (KJM), the Tobias Foundation (KJM), The Magnus Bergvall Foundation (2024-975, AP), the Swedish Research Council (2025-02053, AP) and Science for Life Laboratory (SciLifeLab) (SLL 2026/28, AP). We would like to thank the Flow Cytometry and Genomics Core Facilities at Oslo University Hospital.

---

**Supplemental Figure 1.**
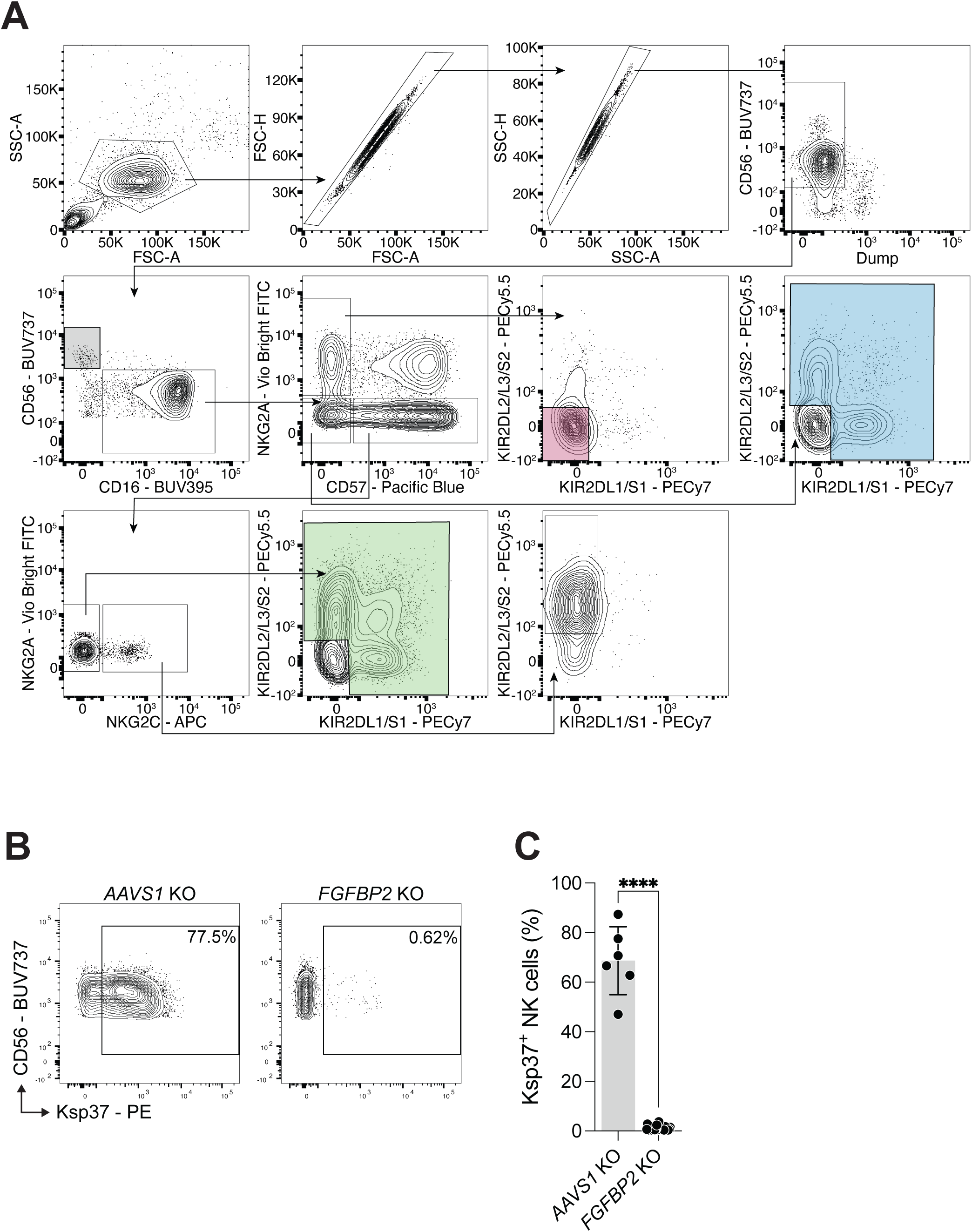
Ksp37 expression in NK cell subsets. (**A**) Gating strategy for NK cell subsets using flow cytometry. (**B**) Representative flow cytometry staining of Ksp37 in *AAVS1* control or *FGFBP2* knock-out (KO) NK cells. (**C**) Percentage of Ksp37^+^ cells in *AAVS1* control or *FGFBP2* KO NK cells as quantified using flow cytometry. Mann-Whitney test was performed in (**C**). Bars show the mean and error bars the standard deviation. *n* = 6 – 36 donors. \*\*\*\**p* < 0.0001.

**Supplemental Figure 2.**
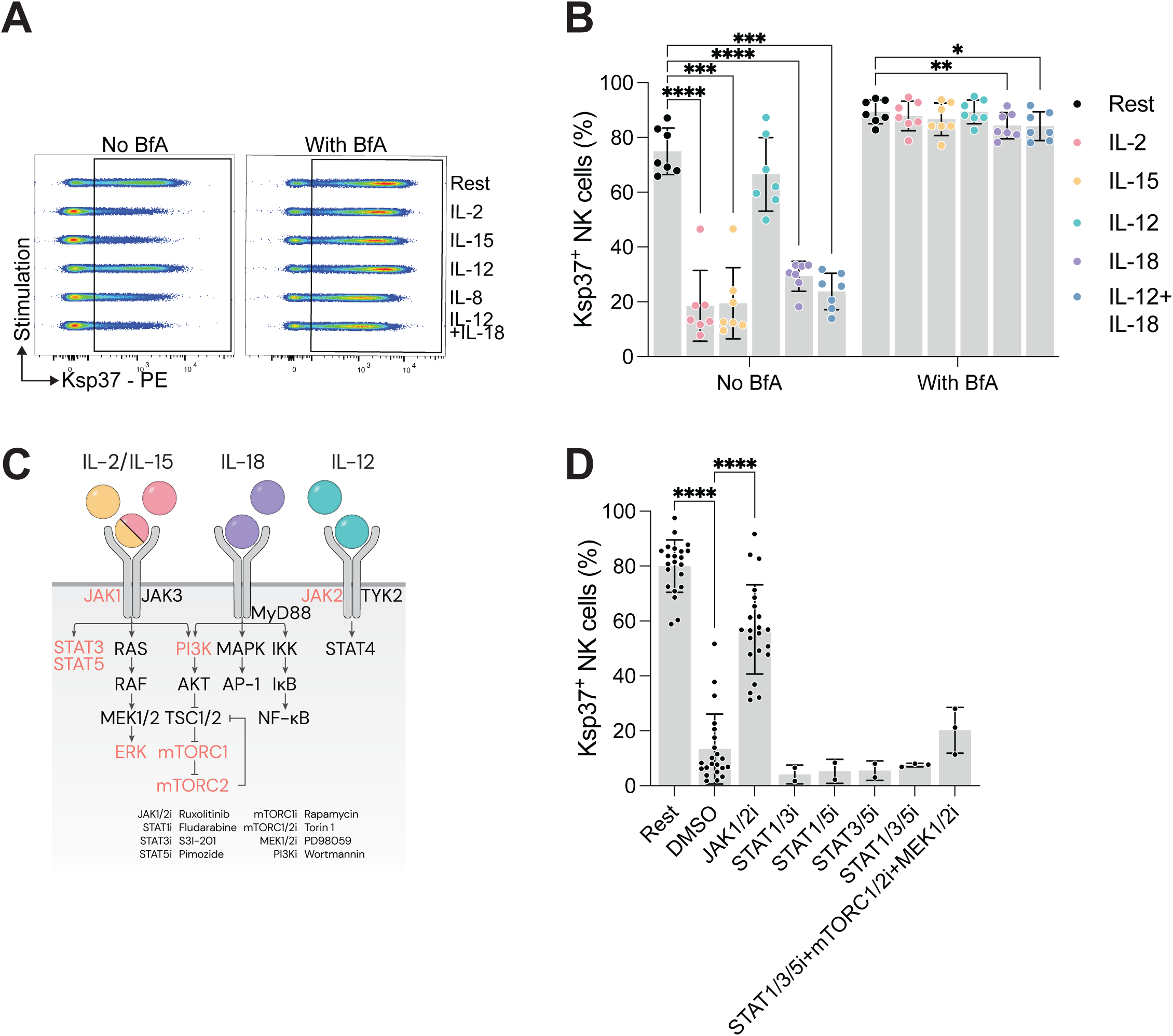
Rapid secretion of Ksp37 upon cytokine stimulation. (**A**) Representative flow cytometry staining showing Ksp37 expression in NK cells cultured overnight with different cytokines in presence or absence of brefeldin A (BfA). (**B**) Percentage of Ksp37^+^ NK cells upon overnight culture with different cytokines in presence or absence of brefeldin A (BfA). (**C**) Schematic of main signaling pathways downstream of stimulation with IL2, IL15, IL12 and IL18. Signaling molecules highlighted in orange are targeted by inhibitors. (**D**) Percentage of Ksp37^+^ NK cells upon rest or with 5 ng/mL IL-15 in presence of indicated signaling pathway inhibitors (i) or DMSO vehicle control. Repeated measures two-way ANOVA with Geisser-Greenhouse corrected followed by Šídák’s multiple testing comparison test was performed in (**B**). Kruskal-Wallis test followed by Dunn’s multiple comparison test was performed in (**D**). Bars show the mean and error bars the standard deviation. *n* = 2 – 22 donors. \*\*\*\**p* < 0.0001; \*\*\**p* < 0.001; \*\**p* < 0.01; and \**p* < 0.05.

**Supplemental Figure 3.**
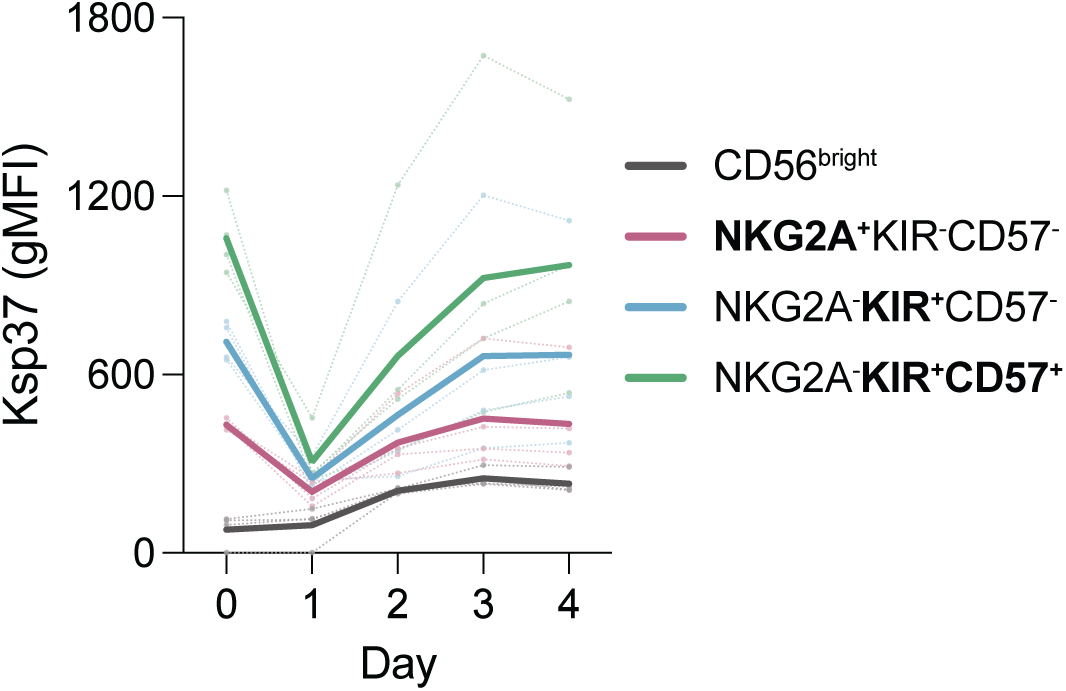
Reloading of Ksp37 in NK cell subsets. Ksp37 gMFI within NK cell subsets before stimulation (day 0) and on days 1-4 of stimulation with 5 ng/mL IL-15. Solid lines indicate the mean and dashed lines indicate individual donors (*n* = 4).

